# Interleukin-6 Regulates the Neutrophil Response to Diverse Bacteria

**DOI:** 10.64898/2026.01.08.698217

**Authors:** Justin M. Owens, Hannah K. Weppner, Aitana Ignes-Romeu, Jacob W. Burleson, Laurel E. Hind

## Abstract

Neutrophils are critical mediators of the innate immune response, and their antimicrobial functions are tightly regulated by a myriad of cytokines. Interleukin-6 (IL-6) is known to be essential for an effective immune response; however, how varying IL-6 concentrations affect the neutrophil response remains poorly understood. Because IL-6 concentrations can vary greatly across different disease states, we investigated the concentration dependent effects of IL-6 on the neutrophil response to diverse bacterial pathogens using an infection-on-a-chip microfluidic device. We found that a high exogenous IL-6 concentration (100 ng/mL) reduced neutrophil extravasation, migration speed, and displacement compared to conditions without exogenous IL-6. In contrast, a lower exogenous IL-6 concentration (10 ng/mL) produced pathogen-specific effects on neutrophil extravasation: exogenous IL-6 increased neutrophil extravasation in response to *Pseudomonas aeruginosa*, did not change in response to *Listeria monocytogenes*, and decreased in response to *Staphylococcus aureus* relative to controls. We then determined the potential endothelial cell contributions to these responses. We found that increasing IL-6 concentration resulted in decreased VE-cadherin expression and that 100 ng/mL exogenous IL-6 resulted in lower ICAM-1 expression than 10 ng/mL exogenous IL-6 in an endothelium exposed to *P. aeruginosa*. Together, these results demonstrate that IL-6 exerts concentration- and pathogen-dependent effects on neutrophil recruitment and migration, supporting a dual role for IL-6 as both pro-inflammatory and anti-inflammatory, with higher IL-6 concentrations resulting in a more anti-inflammatory neutrophil response.

## 1 Introduction

Neutrophils are among the first responders in the innate immune response and are essential for pathogen clearance and host defense (1). Their response is tightly regulated by cytokines which bind to receptors on neutrophils and initiate downstream signaling pathways that regulate neutrophil function (1). Dysregulated cytokine concentrations can disrupt these downstream signaling pathways, leading to neutrophil dysfunction that has been implicated in the pathogenesis of many diseases (1). While cytokine concentration dysregulation is a known contributor to disease pathology, how specific cytokine concentrations contribute to neutrophil dysfunction is largely unknown.

Interleukin-6 (IL-6) is an essential pro-inflammatory cytokine, necessary for an effective defense against pathogens, partially attributed to its role in stimulation of endothelial cells and lymphocytes (2–5). Previous work by our lab and others has shown the presence of IL-6 is essential for an effective neutrophil response and inhibiting IL-6 reduces neutrophil recruitment (6–10). However, the effect of varying IL-6 concentrations on the neutrophil response is largely unknown (11,12). There is a need to better understand how IL-6 concentration affects the neutrophil response as IL-6 concentration can vary from levels of 1-5 pg/mL in homeostatic conditions to over 750 ng/mL in meningococcal septic shock patients. In particular, it is especially important to understand how increased IL-6 levels affect neutrophil function as they are frequently correlated with increased severity in conditions such as rheumatoid arthritis, Crohn’s disease, and sepsis (13–16).

A major limitation in studying concentration-dependent effects of IL-6 on the immune system is a lack of physiologically relevant experimental systems in which primary human neutrophil function can be tracked while controlling local IL-6 concentration. Microphysiological systems that incorporate key elements of the infectious microenvironment including relevant geometries, such as model blood vessels, and live pathogens are critical for these types of studies as our lab and others have shown neutrophils do not effectively respond to bacterial infection without an endothelium, and neutrophils do not extravasate without an infection present (7,17). The inclusion of a model blood vessel is also critical as endothelial cells express tight junction proteins such as VE-cadherin and adhesion receptors including ICAM-1 which govern neutrophil extravasation out of the blood vessel (18). Critically, IL-6 concentration is positively correlated with endothelial cell secretion of proinflammatory chemokines IL-8 and MCP-1 and IL-6 signaling in endothelial cells supports neutrophil transmigration (19,20), suggesting concentration-dependent effects of IL-6 on neutrophil function could be indirectly mediated by the vascular endothelium.

In this study, we used our infection-on-a-chip device that incorporates a collagen extracellular matrix, a three-dimensional blood vessel endothelium lined with primary human endothelial cells, live bacteria, and primary human neutrophils to generate a physiologically relevant infection microenvironment where controlled levels of exogenous IL-6 concentration can be added (7). Our lab previously found neutrophils have pathogen-specific responses, therefore, we investigated IL-6 concentration dependent neutrophil function in response to three clinically relevant bacterial pathogens: *Pseudomonas aeruginosa, Listeria monocytogenes*, and *Staphylococcus aureus* (7,21–23). We found that high initial levels of IL-6 reduced neutrophil extravasation added across bacterial species relative to conditions without IL-6, while intermediate levels of IL-6 can have pathogen-dependent effects on neutrophil extravasation. Additionally, high initial levels of IL-6 reduced neutrophil migration speed and displacement over time. To determine the potential for endothelium-mediated regulation, we quantified the expression of ICAM-1 and VE-cadherin on the endothelial lumen and we observed trends suggesting higher exogenous IL-6 levels may lead to decreased VE-cadherin and ICAM-1 expression in the presence of bacteria. Together, these findings indicate that high levels of IL-6 reduce neutrophil extravasation, speed, and displacement, potentially through changes in VE-cadherin and ICAM-1 expression on endothelial cells.

## 2 Materials and Methods

### 2.1 Experimental Methods

#### 2.1.1 Infection-on-a-Chip Device Fabrication

Microfluidic devices were fabricated as previously described (7,17). In brief, two polydimethylsiloxane (PDMS) layers were patterned from top and bottom masters made of PC-Like Advanced High Temp (Accura 5530) resin with a natural finish for the device (Protolabs Inc), aligned, and then bonded to a glass bottom MatTek dish using oxygen plasma (Diener Electronic Femto Plasma Surface System). 4 mg/mL Type I rat tail collagen (354249, Corning) neutralized to a pH of 7.2 was added to devices and allowed to polymerize at room temperature around a 0.337 mm inner diameter PDMS rod before incubating at 37°C and 5% CO_2_ for at least 2 hours. The PDMS rods were removed, and lumens were seeded with HUVECs at 2.0×10^4^ cells/μL, incubated at 37°C and 5% CO_2_ and flipped every 15 min for 1 hour before performing a media change with 37°C EGM-2. The devices were then cultured for 2 days at 37°C and 5% CO_2_ with twice daily EGM-2 media changes.

#### 2.1.2 Collection of Neutrophils from Human Participants

All blood samples were collected from donors aged 18-65 in accordance with our institutional review board-approved protocol (#20-0082) per the Declaration of Helsinki. Blood was collected in BD Vacutainer EDTA Tubes (366643, BD Biosciences).

#### 2.1.3 Neutrophil Isolation

Primary human neutrophils were isolated from whole blood samples using the MACSxpress Neutrophil Isolation Kit (130-104-434, Miltenyi Biotec) and the Erythrocyte Depletion Kit (130-098-196, Miltenyi Biotec) in accordance with manufacturer protocols.

#### 2.1.4 Bacteria Preparation

*Pseudomonas aeruginosa* (strain K), *Listeria monocytogenes* (serotype 1/2a strain 10403s) and *Staphylococcus aureus* (strain USA300 LAC) were cultured as previously described (7). In brief, ∼16 hours before an experiment, a single colony from a streaked plate is collected using an inoculation loop and put into 5 mL of media (LB broth for *P. aeruginosa*, and Brain Heart Infusion broth for *L. monocytogenes* and *S. aureus*). Approximately 75 min prior to the experiment, 1 mL of this solution is diluted with 4 mL of additional broth. At the time of the experiment, 1 mL of the bacterial solution was spun down at 17,000xg for 1 min and resuspended in 100 μL of EGM-2. A 1:100 diluted solution was created by adding EGM-2 and the OD was measured. The final volume of the bacteria was adjusted so that all bacteria had a final concentration of 1.25×10^6^ CFU/mL which corresponds to OD = 5.0 for P. aeruginosa, OD = 2.3 for *L. monocytogenes*, and OD = 10.0 for *S. aureus*.

#### 2.1.5 Human Umbilical Vein Endothelial Cell (HUVEC) Culture

Pooled human umbilical vein endothelial cells (HUVEC, 50-305-964, PromoCell) cultured in T-75 flasks and maintained in Endothelial Growth Media (EGM-2, NC9525043, Lonza Walkersville CC3162) until they reached 80% confluence. These cells were used from passages 2 to 6.

#### 2.1.6 Confocal Image Acquisition

Confocal images were acquired using a Nikon A1R HD25 Laser Scanning Confocal Microscope with either a Nikon 10X/0.45 (NA) air objective (extravasation, permeability, and viability experiments) or a Nikon 20X/0.75 (NA) air objective (migration and lumen staining experiments) and Nikon Elements acquisition software. Samples were maintained at 37 °C, 5.0% CO_2_, and 90% relative humidity using an OKO Labs environmental chamber.

#### 2.1.7 Neutrophil Extravasation and Migration Experiments

After 2 days of culture, 0, 10, 100 ng/mL IL-6 (130-093-929, Miltenyi Biotec) is added to the lumens and incubated at 37 °C and 5.0% CO_2_ for approximately 45 minutes. At the end of this incubation, approximately 4 μL of EGM-2 containing neutrophils at a concentration of 7.5×10^6^ cells/mL with 50% of the neutrophils stained with calcein AM (C3100MP, Invitrogen) or calcein red-orange (C34851, Invitrogen), were added to each lumen. About 3 μL of EGM-2 with 1.25×10^6^ CFU/mL of a bacterial species was added to the top port of the device, and then devices were imaged every hour for 8 hours for extravasation. Migration images were taken concurrently every 30 seconds for 10 minutes every hour for 8 hours.

#### 2.1.8 Viability Staining

Immediately following an 8-hour extravasation/migration experiment, a solution containing 2 μg/mL of propidium iodide (5370605ML, Sigma) in EGM-2 was added to the side and top ports of the device and incubated for 10 minutes at room temperature before imaging a 400 μm stack in the Z-direction for each lumen, centered at the middle of the lumen.

#### 2.1.9 HUVEC Protein Expression Staining Assay

0 ng/mL, 10 ng/mL, or 100 ng/mL IL-6 in EGM-2 was added to each lumen and incubated at 37°C with 5% CO_2_ for 2 hours. For experiments containing *P. aeruginosa*, about 3 μL of EGM-2 with 1.25×10^6^ CFU/mL of *P. aeruginosa* was placed on the top port of the device and incubated with the lumens containing different IL-6 concentrations during the same 2-hour incubation. Lumens were fixed by adding prewarmed 4% paraformaldehyde (PFA, AAJ19943K2, Thermo Scientific) to the side ports and incubating at 37°C for 30 minutes. PFA was removed with 3 washes with PBS with calcium and magnesium (PBS+, 14040-177, Gibco). To permeabilize, a solution of 0.2% BSA (A7906, Sigma-Aldrich) and 0.1% Tween 20 (E108, Bethyl Laboratories) in PBS+ was added to each device 3 times and incubated at room temperature for 10 minutes. Next, a staining solution of 0.005 mg/mL Hoechst 34580 (H21486, Invitrogen), 5:600 anti-ICAM-1 (FITC conjugated, BBA20, R&D Systems), 1:1000 phalloidin (iFluor-555 conjugated, ab176756, Abcam), and 5:600 anti-VE-Cadherin (Alexa Fluor 647 conjugated, 561567, BD Pharmingen) was added 3 times to each device. Devices were incubated overnight at 4°C and protected from light. Before imaging, devices were rinsed 3 times with PBS+ to remove excess stains.

### 2.2 Image Processing and Data Analysis Methods

#### 2.2.1 Neutrophil Extravasation and Migration Analysis

The images from extravasation experiments were converted to max intensity projections in the Z-axis. An outer region of interest was drawn for every lumen by drawing a rectangular region approximately spanning from the midpoint of the diagonal of the top half of the ECM barrier, down to the lumen edge and across to the other side of the ECM barrier so that the edges of the device are not included in the outer region of interest. An inner region of interest was drawn by matching the horizontal width of the outer region of interest and adjusting the height to be the height of the lumen. The number of neutrophils in each region of interest were counted at each time point using Nikon Elements analysis software. The number of neutrophils in the outer region of interest (extravasated neutrophils) was normalized by the maximum number of neutrophils in the inner region of interest (un-extravasated neutrophils) across all time points for that lumen. Neutrophils that had extravasated were tracked using the Cell Motility function in Nikon Elements. Tracks that corresponded to less than half of the time points for each hour were excluded as they represented neutrophils that moved out of frame.

#### 2.2.2 Neutrophil Viability Analysis

Images were converted to a max intensity projection in the Z-axis and analyzed using a region of interest approximately equal to the sum of the inner and outer regions of interest used in the extravasation analysis. The percentage of viable cells was calculated by dividing the total number of cells stained with calcein (live) by the sum of live cells and cells stained with propidium iodide (dead).

#### 2.2.3 HUVEC Protein Expression Analysis

The raw data from the HUVEC protein expression experiments were transferred to FIJI (ImageJ) for saving individual channels and cropping images to the approximate size of the endothelium. No changes to the brightness/fluorescence intensity were made during this process. These pictures were read using a MATLAB script that segments nuclei based on the fluorescence intensity, area, and eccentricity and measures the fluorescence intensity of the pixels in the VE-cadherin or ICAM-1 images and totals them. The values for total fluorescence intensity were normalized by dividing by the number of nuclei for each lumen. Average protein expression (total fluorescence intensity) was calculated for each biological replicate (lumens with HUVECs from the same flask) for each IL-6 condition (0, 10, 100 ng/mL). The fold change in protein expression was calculated within each biological replicate by dividing each condition’s average protein expression value by the average protein expression value for 0 ng/mL in that biological replicate. The fold change values of all biological replicates were analyzed together to compare the 0, 10, and 100 ng/mL conditions to each other in the absence or presence of *P. aeruginosa*.

#### 2.2.4 Statistical Analysis

For all experiments, data was pooled from three or more independent replicates. For the extravasation, migration and protein expression data, a one-way analysis of variance (ANOVA) was performed in MATLAB/RStudio, followed by Tukey’s Honest Significant Difference (HSD) test of each IL-6 concentration for each time point. The viability data was analyzed in MATLAB using a two-way ANOVA followed by a Tukey’s HSD test. For protein expression analysis, the data was analyzed in MATLAB using a one-way ANOVA followed by a Tukey’s HSD test. Each bar represents the mean plus standard error of the mean (SEM). P values are labeled as * P<0.05, † P<0.05, # P<0.001.

## 3 Results

### 3.1 High IL-6 Levels Limit Neutrophil Extravasation to Diverse Bacteria

The initial step in the neutrophil response to bacterial infections is extravasation across the vascular endothelium into nearby infected tissues. To determine how IL-6 concentration regulates neutrophil extravasation out of a model blood vessel, we incubated the endothelium of our infection-on-a-chip device with IL-6 at 0, 10, or 100 ng/mL for approximately 45 min, followed by the addition of neutrophils in media containing the corresponding IL-6 concentration. Neutrophil extravasation was quantified over 8 hours in response to one of three bacterial species (*P. aeruginosa, L. monocytogenes*, or *S. aureus*) (Figure 1A,C,E). Consistent with our prior study, we observed the highest extravasation levels in response to *L. monocytogenes* and the lowest extravasation levels in response to *S. aureus* at the 0 ng/mL condition (7). Interestingly, the addition of IL-6 altered extravasation in a concentration- and pathogen-dependent manner. In response to *P. aeruginosa*, 10 ng/mL exogenous IL-6 significantly increased extravasation relative to the 100 ng/mL exogenous IL-6 at the 2-, 4-, and 6-hour timepoints (P<0.001) and 0 ng/mL exogenous IL-6 at the 2-hour timepoint (P<0.05) (Figure 1B). Additionally, 100 ng/mL exogenous IL-6 reduced extravasation relative to 0 ng/mL exogenous IL-6 at the 6-hour timepoint (P<0.05). In contrast, neutrophils responding to *L. monocytogenes* did not display statistically significant differences in extravasation across the IL-6 concentrations, but a consistent trend toward lower extravasation at 100 ng/mL exogenous IL-6 was found, as with *P. aeruginosa* (Figure 1D). *S. aureus* also elicited IL-6 concentration-dependent neutrophil extravasation, with significantly reduced extravasation for 10ng/mL exogenous IL-6 at 4 and 8 hours and 100 ng/mL at 2, 4 and 8 hours, relative to 0 ng/mL exogenous IL-6 (Figure 1F). To determine if neutrophil viability was affecting extravasation levels, cells in the microfluidic device were stained with propidium iodide following an 8-hour extravasation experiment. We found neutrophil viability remained >97.5% at all IL-6 concentrations in response to *P. aeruginosa* with no significant change between IL-6 conditions (Figure S1). Overall, these results indicate that IL-6 concentration modulates neutrophil extravasation out of the lumen in a pathogen-specific manner and the observed differences are not due to neutrophil death.

**Figure 1:**
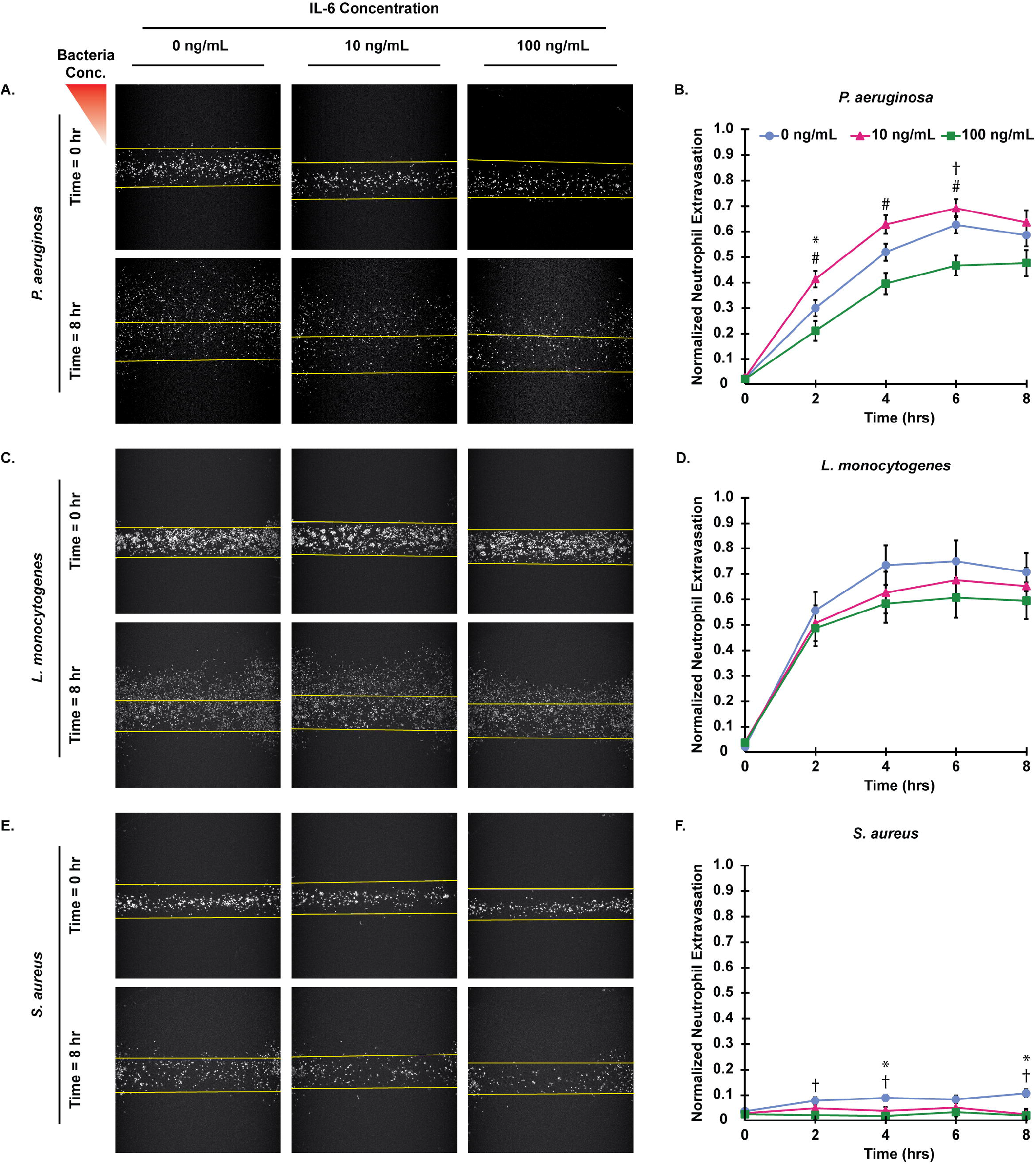
IL-6 concentration affects neutrophil extravasation in a pathogen-specific manner. (A,C,E) Representative images of extravasated neutrophils preincubated with 0, 10, or 100 ng/mL IL-6 at 0 and 8 hours post stimulation with (A) *P. aeruginosa*, (C) *L. monocytogenes*, or (E) *S. aureus* (scale bar = 250 μm). Yellow lines indicate the edge of the lumen. (B, D, F) Normalized number of extravasated neutrophils in microfluidic devices preincubated with 0, 10, or 100 ng/mL IL-6 responding to (B) *P. aeruginosa*, (D) *L. monocytogenes*, or (F) *S. aureus*. Data quantified from 14 devices for 0 ng/mL, 13 devices for 10 ng/mL, and 10 devices for 100 ng/mL across 4 independent experiments and 3 neutrophil donors for *P. aeruginosa*. Data quantified from 15 devices for 0 ng/mL, 13 devices for 10 ng/mL, and 13 devices for 100 ng/mL across 4 independent experiments and 3 neutrophil donors for *S. aureus*. Data quantified from 11 devices for 0 ng/mL, 13 devices for 10 ng/mL, and 12 devices for 100 ng/mL across 4 independent experiments and 3 neutrophil donors for *L. monocytogenes*. All IL-6 conditions were compared to each other at each time point. Conditions at each time point were compared with a one-way ANOVA and then Tukey’s Honestly Significant Difference (HSD) post-hoc test was used to calculate p-values. Error bars indicate the mean ± SEM. * indicates significance between the 0 and 10 ng/mL conditions at the same time point where p<0.05. † indicates significance between the 0 and 100 ng/mL conditions at the same time point where p<0.05. # indicates significance between the 10 and 100 ng/mL conditions at the same time point where p<0.001.

### 3.2 Extended Exposure to High IL-6 Levels Hinders Neutrophil Migration

Following extravasation, neutrophils are directed by inflammatory signals from the endothelium, including IL-6, and bacteria to migrate through the tissue to the source of infection. To determine how IL-6 concentration affects neutrophil migratory properties, extravasated neutrophils were imaged every 30 seconds for 10 min every hour and neutrophil migration speed and displacement were quantified (Figure 2A). Neutrophils responding to *P. aeruginosa*, had significantly reduced migration speed with 100 ng/mL exogenous IL-6 relative to 0 ng/mL exogenous IL-6 over time and reduced displacement with 100 ng/mL relative to the 0 ng/mL and 10 ng/mL exogenous IL-6 at later time points (Figure 2B,C). Neutrophils responding to *L. monocytogenes*, had significantly reduced speed and displacement with 10 ng/mL and 100 ng/mL exogenous IL-6 conditions compared to 0 ng/mL exogenous IL-6 at later timepoints (Figure 2D,E). Migration data for *S. aureus* were not analyzed due to the low number of extravasated neutrophils, which prevented reliable tracking and statistical comparison. Together, these results indicate that high IL-6 concentrations reduce neutrophil migration speed and displacement, and this trend is consistent in neutrophil responses to *P. aeruginosa* and *L. monocytogenes*.

**Figure 2:**
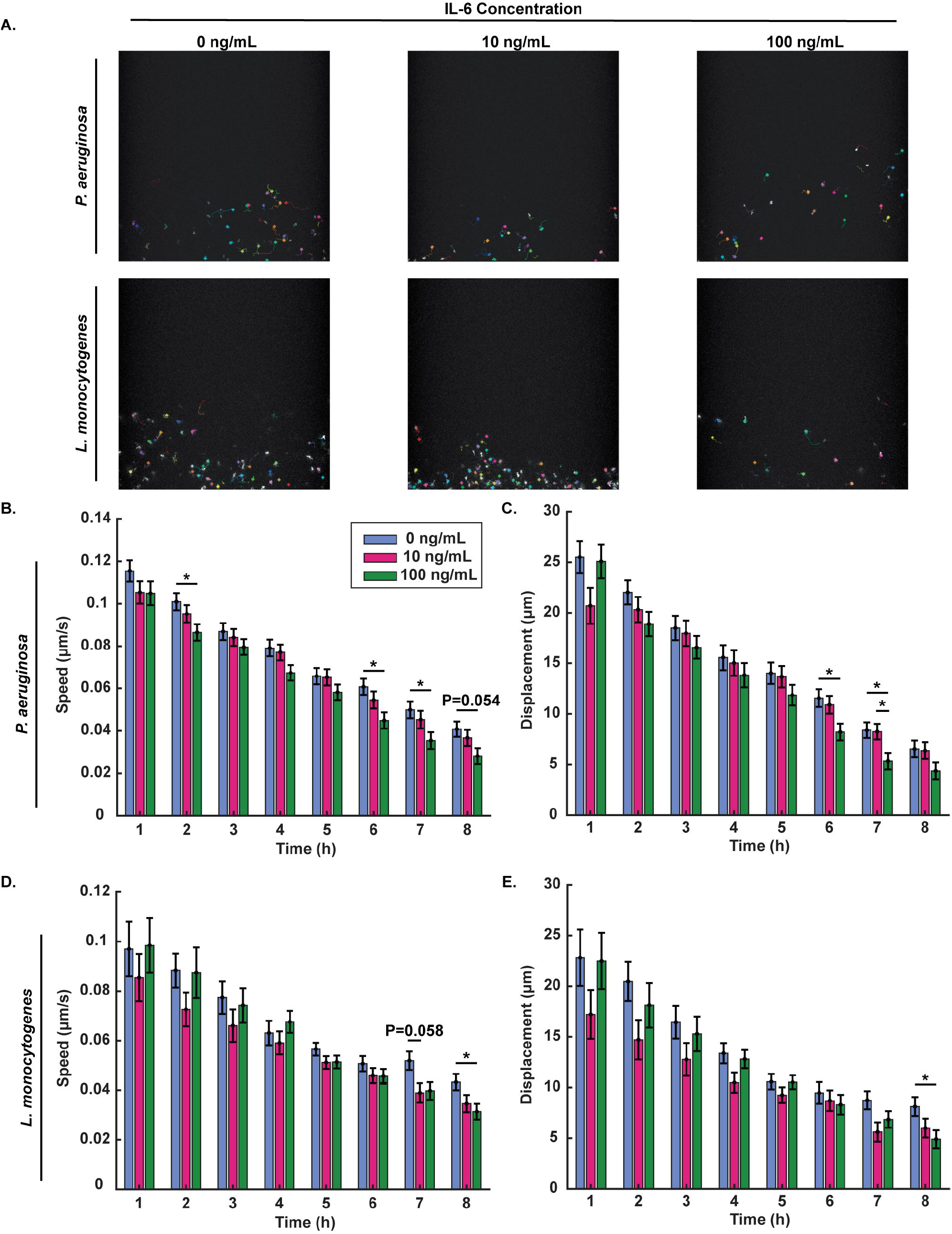
Higher IL-6 levels decrease neutrophil speed and displacement at later timepoints. (A) Representative images and tracks of migrating neutrophils with 0, 10 or 100 ng/mL IL-6 at the 4-hour timepoint after stimulation with *P. aeruginosa* or *L. monocytogenes* (scale bar = 250 μm). Each individual track is shown using different colors. Yellow lines indicate the lumen edge. (B) Neutrophil migration speed and (C) displacement of neutrophils quantified over 10-minute intervals every hour for 8 hours after stimulation with *P. aeruginosa*. (D) Neutrophil migration speed and (E) displacement of neutrophils quantified over 10-minute intervals every hour for 8 hours after stimulation with *L. monocytogenes*. Data quantified from 11 devices for 0 ng/mL, 10 devices for 10 ng/mL, and 11 devices for 100 ng/mL across 3 independent experiments and 3 neutrophil donors for *P. aeruginosa*. Data quantified from 10 devices for 0 ng/mL, 10 devices for 10 ng/mL, and 11 devices for 100 ng/mL across 3 independent experiments and 3 neutrophil donors for *L. monocytogenes*. Cells were tracked using Cell Motility function in Nikon Elements. All IL-6 conditions were compared to each other at each time point. Conditions at each time point were compared using a one-way ANOVA and then p-values were determined using Tukey’s HSD post-hoc test. Error bars indicate the mean ± SEM. Asterisks indicate significance between conditions at a given timepoint (*p<0.05).

### 3.3 Increasing IL-6 Concentration May Decrease VE-Cadherin Expression

To determine whether IL-6-dependent changes in neutrophil extravasation were indirectly due to changes in endothelial junctional protein expression, endothelial cells in our device were incubated with IL-6 and stained for the junctional protein VE-cadherin (Figure 3A,C; unprocessed images shown in Figure S2A). In the absence of bacteria, we found that VE-cadherin expression was unaffected by IL-6 concentration after 2 hours (Figure 3B). In contrast, in the presence of *P. aeruginosa*, VE-cadherin expression was reduced by incubation with 100 ng/mL exogenous IL-6, and, to a lesser extent, by 10 ng/mL exogenous IL-6 (Figure 3D). Together, these findings indicate VE-cadherin is unaffected by exogenous IL-6 when bacteria is not present, but in the context of bacterial infection, increasing IL-6 concentrations are associated with reduced VE-cadherin expression.

**Figure 3:**
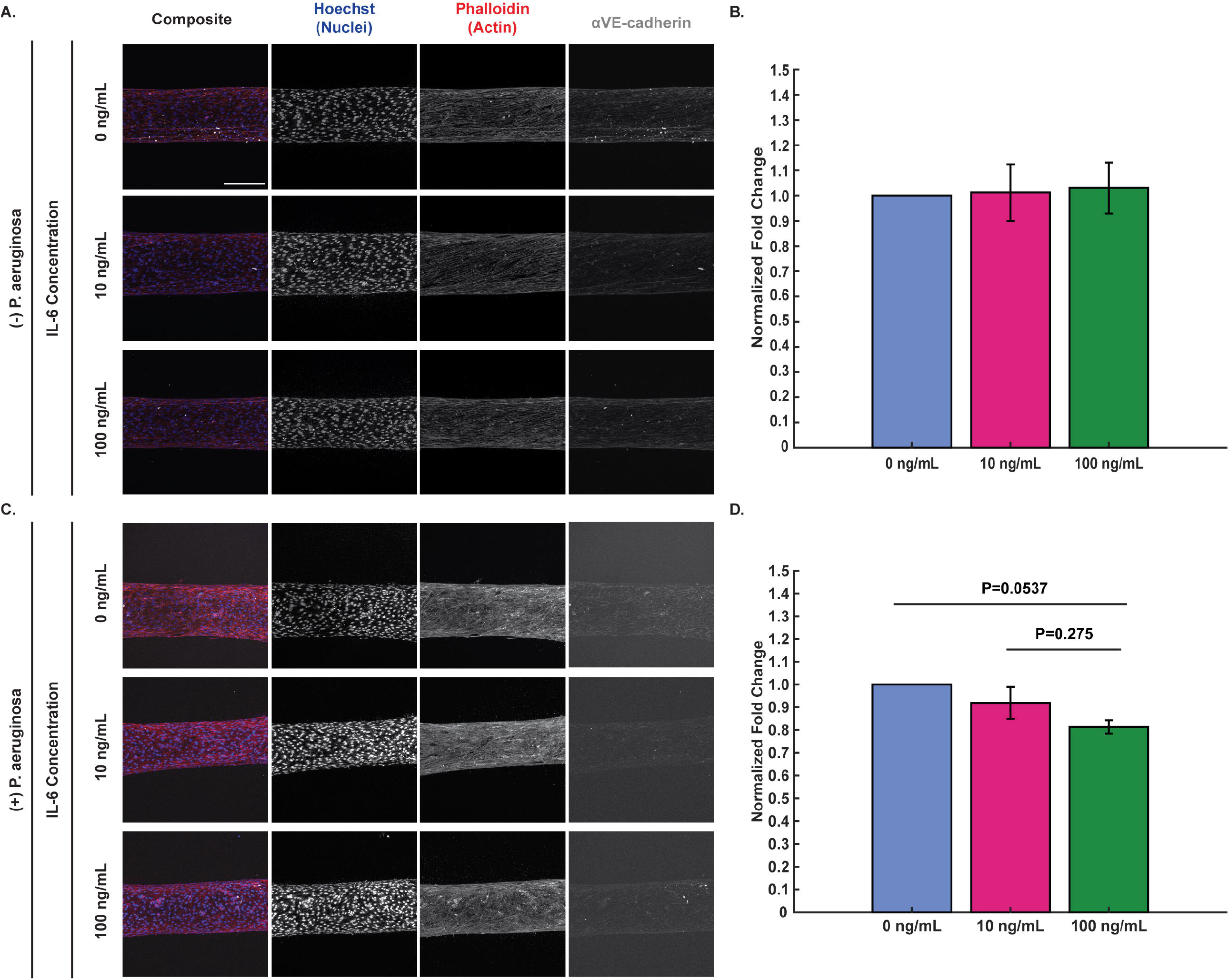
VE-cadherin expression may decrease in presence of exogenous IL-6 and *P. aeruginosa*. Representative maximum intensity projections of confocal images of HUVECs seeded in microfluidic devices and incubated with 0, 10, and 100 ng/mL IL-6 for 2 hours (A) with no bacteria and (C) with *P. aeruginosa*. Cells were fixed and stained with Hoechst (nuclei, blue), phalloidin (actin, red), and anti-VE-cadherin (tight junctions, gray) (scale bar = 250 μm). Images were thresholded in ImageJ to visualize the differences in VE-cadherin. Representative raw images used for protein expression analysis can be found in Figure S2. (B, D) Fold change of VE-cadherin expression was calculated by dividing the total fluorescence intensity of un-thresholded images for each condition by the total fluorescence intensity for the 0 ng/mL IL-6 condition of the corresponding biological replicate. Fold change values for each biological replicate were averaged together and then all IL-6 conditions were compared to each other at each time point. Conditions at each time point were compared using a one-way ANOVA and then p-values were determined using Tukey’s HSD post-hoc test. Error bars indicate the mean ± SEM. (B) Data quantified from 9 devices for 0 ng/mL, 8 devices for 10 ng/mL, and 7 devices for 100 ng/mL across 4 independent experiments for protein expression when not exposed to bacteria. (D) Data quantified from 8 devices for 0 ng/mL, 9 devices for 10 ng/mL, and 8 devices for 100 ng/mL across 3 independent experiments for protein expression when exposed to *P. aeruginosa*.

### 3.4 Increasing IL-6 Concentrations May Decrease ICAM-1 Expression When *P. aeruginosa* is Present

In conjunction with staining for VE-cadherin, endothelial ICAM-1 expression was quantified to understand how IL-6 concentration affects adhesion protein expression and whether this could contribute to the observed changes in neutrophil extravasation (Figure 4A,C; unprocessed images shown in Figure S2B). In the absence of bacteria, exogenous IL-6 elicited a trend of increased ICAM-1 expression compared to 0 ng/mL IL-6 control after 2 hours (Figure 4B). Interestingly, when *P. aeruginosa* was present, incubation with 100 ng/mL exogenous IL-6 decreased ICAM-1 expression relative to 0 and 10 ng/mL exogenous IL-6, with difference between 10 and 100 ng/mL approaching statistical significance (Figure 4D). This reduction in ICAM-1 expression correlates with the decrease in extravasation seen between 0 and 100 ng/mL exogenous IL-6 and between 10 and 100 ng/mL exogenous IL-6 conditions when responding to *P. aeruginosa* (Figure 1B). Together, these results show that exogenous IL-6 increases ICAM-1 expression in endothelial cells when no bacteria is present, and that 100 ng/mL exogenous IL-6 decreases ICAM-1 expression in the presence of *P. aeruginosa* compared to 0 and 10 ng/mL exogenous IL-6.

**Figure 4:**
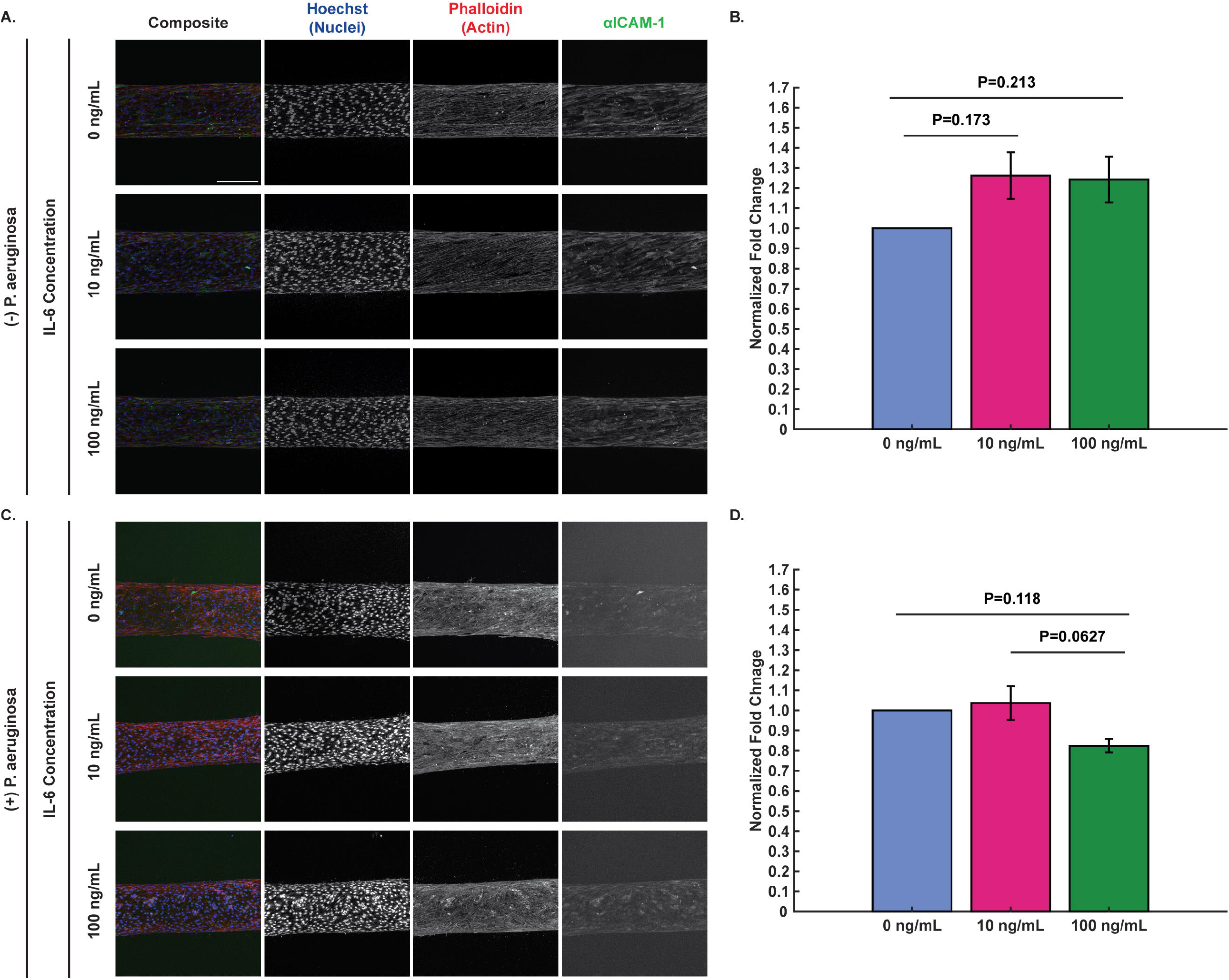
IL-6 increases ICAM-1 expression with no bacteria present, but ICAM-1 expression may decrease in presence of high exogenous IL-6 levels and *P. aeruginosa*. Representative maximum intensity projections of confocal images of HUVECs seeded in microfluidic devices and incubated with 0, 10, and 100 ng/mL IL-6 for 2 hours (A) with no bacteria and (C) with *P. aeruginosa*. Cells were fixed and stained with Hoechst (nuclei, blue), phalloidin (actin, red), and anti-ICAM-1 (neutrophil tight binding to endothelial cells, green) (scale bar = 250 μm). Images were thresholded in ImageJ to visualize the differences in ICAM-1. Representative raw images used for protein expression analysis can be found in Figure S2. (B, D) Fold change of ICAM-1 expression was calculated by dividing the total fluorescence intensity of un-thresholded images for each condition by the total fluorescence intensity for the 0 ng/mL IL-6 condition of the corresponding biological replicate. Fold change values for each biological replicate were averaged together and then all IL-6 conditions were compared to each other at each time point. Conditions at each time point were compared using a one-way ANOVA and then p-values were determined using Tukey’s HSD post-hoc test. Error bars indicate the mean ± SEM. (B) Data quantified from 9 devices for 0 ng/mL, 8 devices for 10 ng/mL, and 7 devices for 100 ng/mL across 4 independent experiments for protein expression when not exposed to bacteria. (D) Data quantified from 8 devices for 0 ng/mL, 9 devices for 10 ng/mL, and 8 devices for 100 ng/mL across 3 independent experiments for protein expression when exposed to *P. aeruginosa*.

## 4 Discussion

In this study, we investigated the effect of IL-6 concentration on the neutrophil response to diverse bacterial species within a physiologically relevant infection-on-a-chip model. We found that exogenous addition of 100 ng/mL IL-6 attenuated the neutrophil responses as evidenced by reduced neutrophil extravasation, as well as migration speed and displacement at later time points. Interestingly, intermediate concentrations of IL-6 had pathogen-dependent effects on the neutrophil response, enhancing neutrophil extravasation in response to *P. aeruginosa*, having no effect in response to *L. monocytogenes*, and reducing extravasation in response to *S. aureus*. Notably, we also found endothelial protein expression of VE-cadherin and ICAM-1 may be indirectly influencing neutrophil function.

We demonstrated, in the absence of bacteria, VE-cadherin expression was unchanged by the different exogenous IL-6 concentrations and ICAM-1 expression increased in the presence of exogenous IL-6, consistent with prior literature (24–27). However, in the presence of *P. aeruginosa*, increasing IL-6 concentration led to decreased VE-cadherin expression. VE-cadherin expression has previously been reported to decrease in response to bacterial products including lipopolysaccharide and LasB toxin, which *P. aeruginosa* secretes; however, to our knowledge, an IL-6 concentration-dependent decrease in VE-cadherin in response to infection has not been reported (28,29). Interestingly, ICAM-1 expression was reduced at 100 ng/mL exogenous IL-6 compared to 10 ng/mL and 0 ng/mL exogenous IL-6 and the comparison between 10 and 100 ng/mL exogenous IL-6 trended towards statistical significance. These changes in ICAM-1 expression match the reduced neutrophil extravasation for 100 ng/mL exogenous IL-6 compared to 0 and 10 ng/mL exogenous IL-6 in response to *P. aeruginosa*.

Our results suggest that high concentrations of IL-6 may result in an anti-inflammatory immune response characterized by reduced neutrophil extravasation and migration. While IL-6 is traditionally thought of as a pro-inflammatory cytokine, there is evidence that suggests IL-6 can cause anti-inflammatory responses. IL-6 signals through two distinct pathways: the classical IL-6 signaling pathway via membrane bound IL-6 receptor (IL-6R) which is pro-inflammatory and occurs on neutrophils and the IL-6 trans-signaling pathway via soluble IL-6R (sIL-6R) which is anti-inflammatory and occurs on all cell types including endothelial cells (4,30). In the context of the neutrophil response, high levels of IL-6 trans-signaling can be thought of as anti-inflammatory because it represents a transition point of the innate immune response where more monocytic cells are recruited instead of neutrophils which is accomplished by downregulating IL-8 and upregulating MCP (31,32). Increased IL-6 levels lead to IL-6R shedding, increasing IL-6 trans-signaling through sIL-6R and shifting the neutrophil response from pro-inflammatory classical signaling to anti-inflammatory trans-signaling (4,30). There is clinical evidence for IL-6 signaling to shift from pro-inflammatory to anti-inflammatory due to increased sIL-6R in human patients with early versus late acute renal allograft rejection (33). Independent of the signaling pathway, IL-6 exerts anti-inflammatory effects by inducing the secretion of IL-1Ra and soluble TNF receptor p55, anti-inflammatory proteins that block the binding of the major pro-inflammatory cytokines IL-1β and TNF-α (34). Either a shift from classical IL-6 signaling to IL-6 trans-signaling or the induction of IL-1Ra or soluble TNF receptor p55 signaling could be causing the anti-inflammatory response observed at high exogenous IL-6 concentration in our device. Future studies investigating sIL-6R levels at different IL-6 concentrations will be critical in establishing a more concrete mechanism for these observations.

IL-6 is an essential regulator of effective immune responses. Despite its importance, very little research has been conducted to understand IL-6 concentration-dependent effects on the immune response. In this study, we demonstrated IL-6 concentration influences whether neutrophil responses are more pro- or anti-inflammatory and that these effects are pathogen-dependent. Future studies into how IL-6 concentrations holistically affect the immune response to diverse bacteria could help create broadly impactful treatments that control IL-6 levels to produce an optimal immune response to various pathogens.

## 6 Author Contributions

J.M.O. and L.E.H designed the research and wrote the manuscript; J.M.O., H.K.W, and J.W.B. performed experiments; J.M.O., H.K.W, A.I.R, and J.W.B. analyzed and interpreted data.

## 8 Acknowledgments

The authors would like to thank the members of the Hind Lab, not listed as authors, for valuable discussions and insights that contributed to this work.

## 9 Data Availability Statement

The raw data supporting the conclusions of this article will be made available by the authors upon request, without undue reservation.

