## Supplemental Data for "Interleukin-6 Regulates the Neutrophil Response to Diverse Bacteria"

Supplementary Material

### 1 Supplementary Figures and Tables

#### 1.1 Supplementary Figure 1

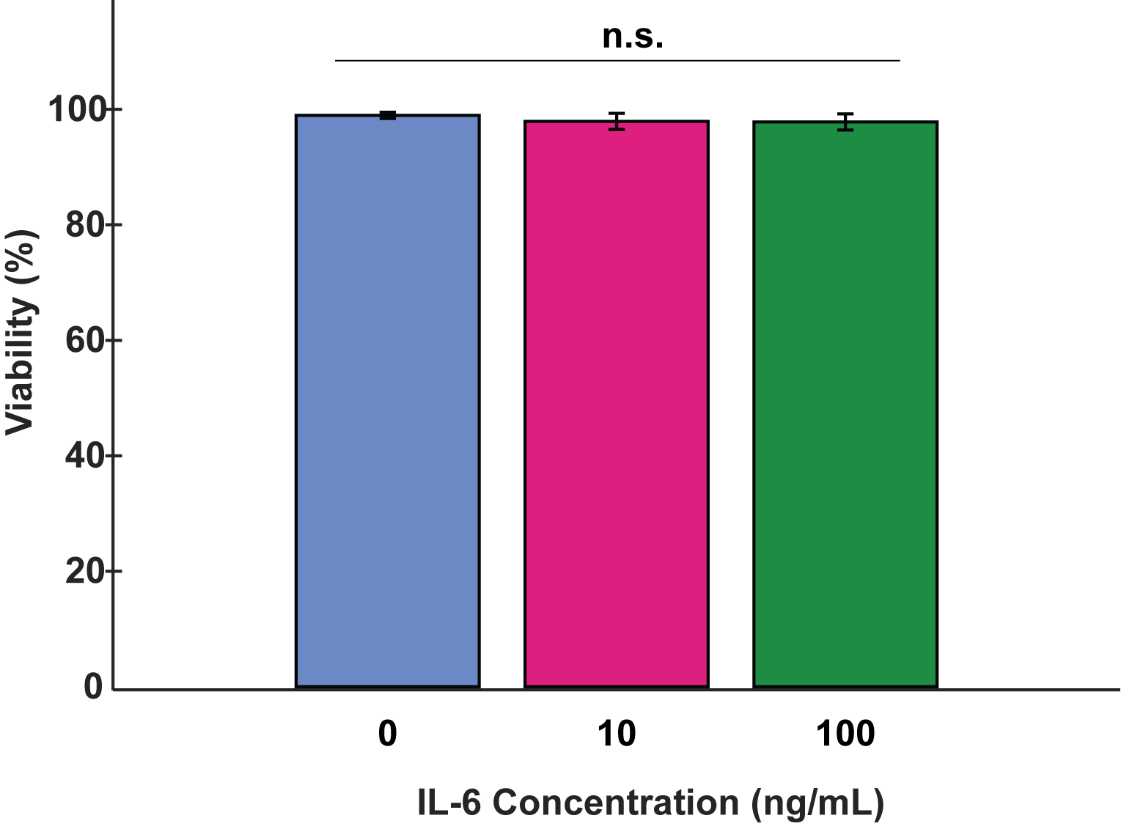

**Figure S1:** **Neutrophil viability unaffected by addition of IL-6**. Propidium iodide was added to microfluidic devices upon the conclusion of an 8-hour extravasation or migration experiment stimulated with *P. aeruginosa*. The percentage of viable cells was calculated by dividing the total number of cells stained with calcein (live) by the sum of live cells and cells stained with propidium iodide (dead). Data quantified from 28 devices across 3 independent experiments and 3 neutrophil donors. All IL-6 conditions were compared to each other at each time point using a two-way ANOVA followed by Tukey’s HSD post-hoc test to determine p-values. Error bars indicate the mean ± SEM.

#### 1.2 Supplementary Figure 2

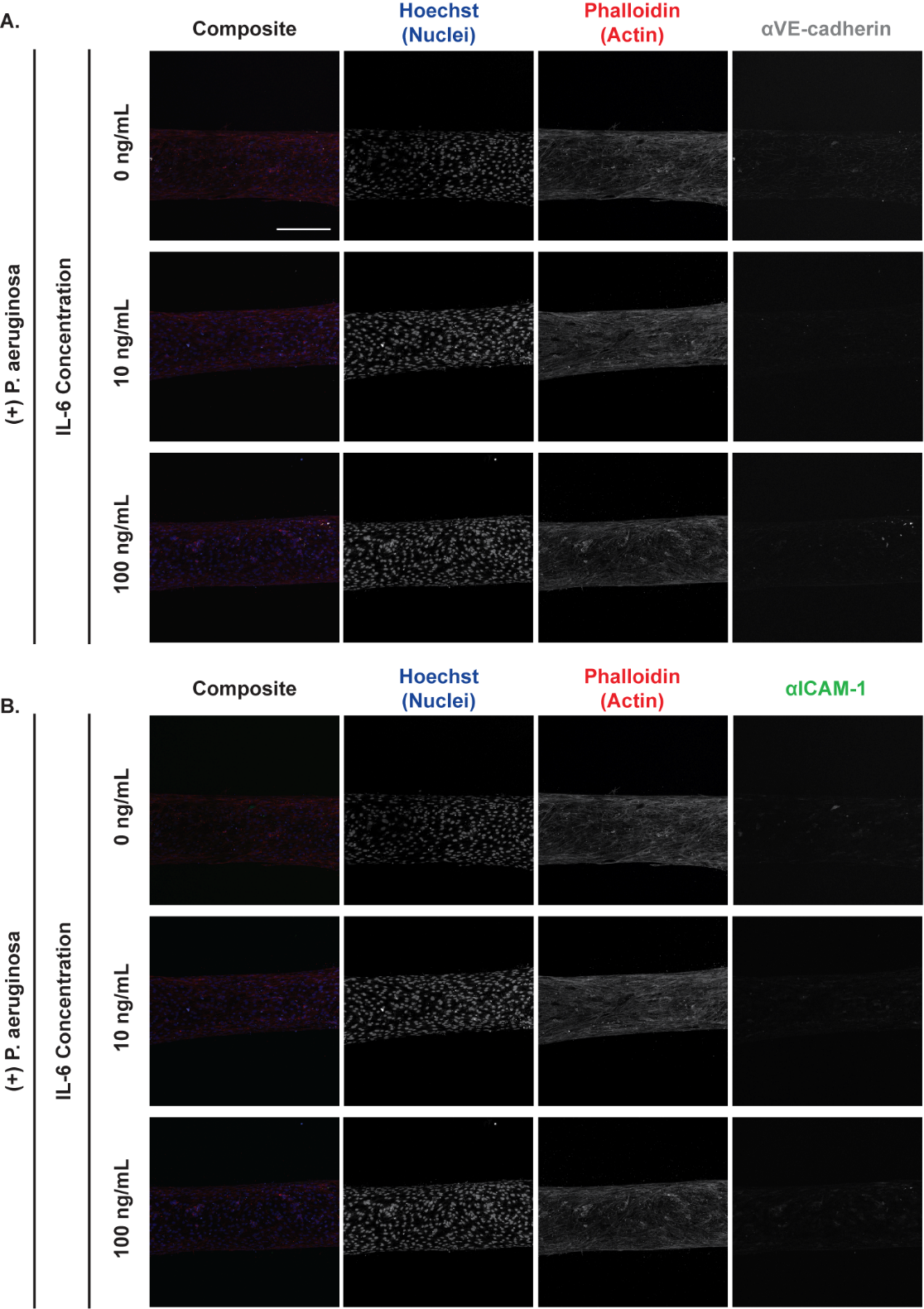

**Figure S2: Unmodified images of VE-cadherin and ICAM-1 fluorescence intensity for conditions with *P. aeruginosa* and exogenous IL-6.** Original maximum intensity projections of confocal images of HUVECs without the additional thresholding steps that were used to brighten the (A) VE-cadherin and (B) ICAM-1 fluorescence. Cells were seeded in microfluidic devices and incubated with 0, 10, and 100 ng/mL IL-6 for 2 hours with *P. aeruginosa*. Cells were fixed and stained with Hoechst (nuclei, blue), phalloidin (actin, red), anti-VE-cadherin (tight junctions, gray), and anti-ICAM-1 (neutrophil tight binding to endothelial cells, green) (scale bar = 250 μm).
